# A fully adapted headstage for electrophysiological experiments with custom and scalable electrode arrays to record widely distributed brain regions

**DOI:** 10.1101/2021.04.06.438732

**Authors:** Flávio Afonso Gonçalves Mourão, Leonardo de Oliveira Guarnieri, Paulo Aparecido Amaral Júnior, Vinícius Rezende Carvalho, Eduardo Mazoni Andrade Marçal Mendes, Márcio Flávio Dutra Moraes

## Abstract

Electrophysiological recordings lead amongst the techniques that aim to investigate the dynamics of neural activity sampled from large neural ensembles. However, the financial costs associated with the state-of-the-art technology used to manufacture probes and multi-channel recording systems make these experiments virtually inaccessible to small laboratories, especially if located in developing countries. Here, we describe a new method for implanting several tungsten electrode arrays, widely distributed over the brain. Moreover, we designed a headstage system, using the Intan^®^ RHD2000 chipset, associated with a connector (replacing the expensive commercial Omnetics connector), that allows the usage of disposable and cheap cranial implants. Our results showed high-quality multichannel recording in freely moving animals (detecting local field, evoked responses and unit activities) and robust mechanical connections ensuring long-term continuous recordings. Our project represents an open source and inexpensive alternative to develop customized extracellular records from multiple brain regions.

## INTRODUCTION

In 1957, the Nobel laureates David Hunter Hubel and Torsten Nils Wiesel developed sharpened and vinyl insulated tungsten microelectrodes to record the extracellular action potentials in the primary visual cortex of anesthetized and unrestrained cats. (Hubel, 1957; Hubel and Wiesel, 1962). Beside the scientific breakthrough regarding visual system physiology (i.e. the complex cortical representation of visual information), the tungsten microelectrodes, with tip diameter less than a micron, formed the basis for the development of modern neural probes (Hong and Lieber, 2019; Szostak et al., 2017).

Since Hubel and Wiesel’s discovery, electrophysiologists have bolstered creativity-guided hypotheses to design specialized electrode arrays aimed at recording not just a small number of single units but several neuronal populations at once (Hong and Lieber, 2019; Szostak et al., 2017). By making use of metal (Dowben and Rose, 1953; Green, 1958; Hubel, 1957; McNaughton et al., 1983) or silicon (Buzsáki et al., 2015; Jones et al., 1992; Jun et al., 2017; Rousche and Normann, 1998) a wide range of electrodes with different sizes, shapes, and geometries, has been used to sample brain activity. Moreover, alongside the development of specialized electrodes, electrophysiology data acquisition systems have reached a high degree of accuracy, miniaturization and data processing capabilities that allow recording from several channels simultaneously. Nowadays, with the low noise integrated amplifier chips, that digitize signals on the spot (e.g., Intan Technologies - Harrison RR, 2008), researchers are able to record thousands of neurons in a single multi-channel probe (Jun et al., 2017) while transmitting the digitized data from each channel through a very small number of wires.

However, the main problem of these state-of-the-art tools is the high costs that make it impossible to perform cutting-edge experiments, especially in newly-formed under budget laboratories, or in developing countries with a lack of financial resources (Hallal, 2021). Furthermore, in the case of manufactured electrode arrays, the high costs are associated with manufacturer limitations such as the number of brain substrates that can be recorded simultaneously, the headstage sizes, and the need for special connectors and cables that drive up costs even more (Chung et al., 2019; Jun et al., 2017; Voigts et al., 2013).

To successfully deal with the costs and in an attempt to create more versatile electrode arrays, some authors have developed creative and custom-made solutions for recording the Local Field Potential (LFP) and the multiunit activity in freely moving animals (França et al., 2020; Polo-Castillo et al., 2019). Nevertheless, these solutions still need expensive commercial connectors, which are not easily recoverable after the experiment, present a reasonably rigid geometry that does not allow recording widely distributed brain regions and, often have mechanical problems if long-term continuous recordings are intended.

In this work, we describe a full low-cost headstage for electrophysiology experiments with scalable electrode arrays, custom-made to the experimental design. In other words, we present a simple and new architecture for electrode placement, so that a different number of electrode bundles can be widely distributed over the brain. In addition, in order to provide more affordable costs, we propose an adaptation to the RHD2000 headstage system (Intan Technologies^®^), associated with a different connector, to replace the commercial costly ones commonly used in the scientific community (Omnetics Connector Corporation).

## MATERIAL AND METHODS

### Overview

Considering that the main objective of this work is to develop an inexpensive tool to perform electrophysiology recordings, we designed our own printed circuit boards and purchase materials from bulk suppliers, except the integrated amplifier chip developed by Intan Technologies (Harrison RR, 2008) and the data acquisition system from Open Ephys (Siegle et al., 2017). Our design is fully compatible with Open Ephys open-source solution available to the scientific community. It is worth mentioning that the Intan chip performs the most demanding tasks of data acquisition right at the recording site, that is, it amplifies, multiplexes, digitizes each channel with 16-bit resolution, and measures impedance (Harrison RR, 2008).

Our solution is composed by three main parts: 1) headstage with RHD2000 chipset and a SMD/FPC (Flexible Printed Circuit) female connector; 2) a SMD/FPC compatible flat shaped printed circuit board (to be implanted during surgery) with a grid of through holes for soldering electrodes; 3) a fiber board template with drilled stereotaxic coordinates where the electrodes are aligned and fixed (Figure1). Moreover, extra adapters were developed to convert Omnetics to the SMD/FPC connector (in order to allow the use of previous comercial solutions), and generic tools were made via 3D printing to facilitate the manufacture and handling of the electrode arrays (Supplementary material; https://cutt.ly/UcSeYPd).

### Recording Headstage

The recording headstage, although essentially similar to the Intan Technologies standard (https://intantech.com/RHD_headstages.html), has five important changes: 1) The circuit was built in a horizontal setup, significantly reducing damage due to excessive torque induced by sudden movements, long term continuous recordings or seizures. In addition, the horizontal setup facilitated animal movement during the behavioral tasks; 2) With a short circuit in the ‘R2’ pad, or just left empty, RHD 2216 chip (16-channel with differential inputs) or RHD 2132 chip (32-channel with unipolar inputs and common reference) can be soldered to the same PCB; 3) With a short circuit in the ‘R1’ pad and ‘R3’ pad empty, or a short circuit in the ‘R3’ pad and ‘R1’ pad empty, the two types of SPI communication, single ended or low voltage differential signaling (LVDS) can be respectively chosen; 4) The standard Omnetics connector (i.e. model: A79042 or A79046) has been replaced with a SMD/FPC (Flexible Printed Circuits) connector (MKTechnic - model: FPC0520VH-20P. VERTICAL, TYPE 0.5MM, PITCH 2MM), which reduced costs significantly and can be easily replaced in case of damage; 5) The standard Omnetics connector (PZN-12 polarized nano connectors), whose output was intended for the SPI communication, has been exchanged for through holes (pitch - 2 mm) which makes possible to solder either pin bar connectors or wires directly (Figure 1B).

**Figure 1.**
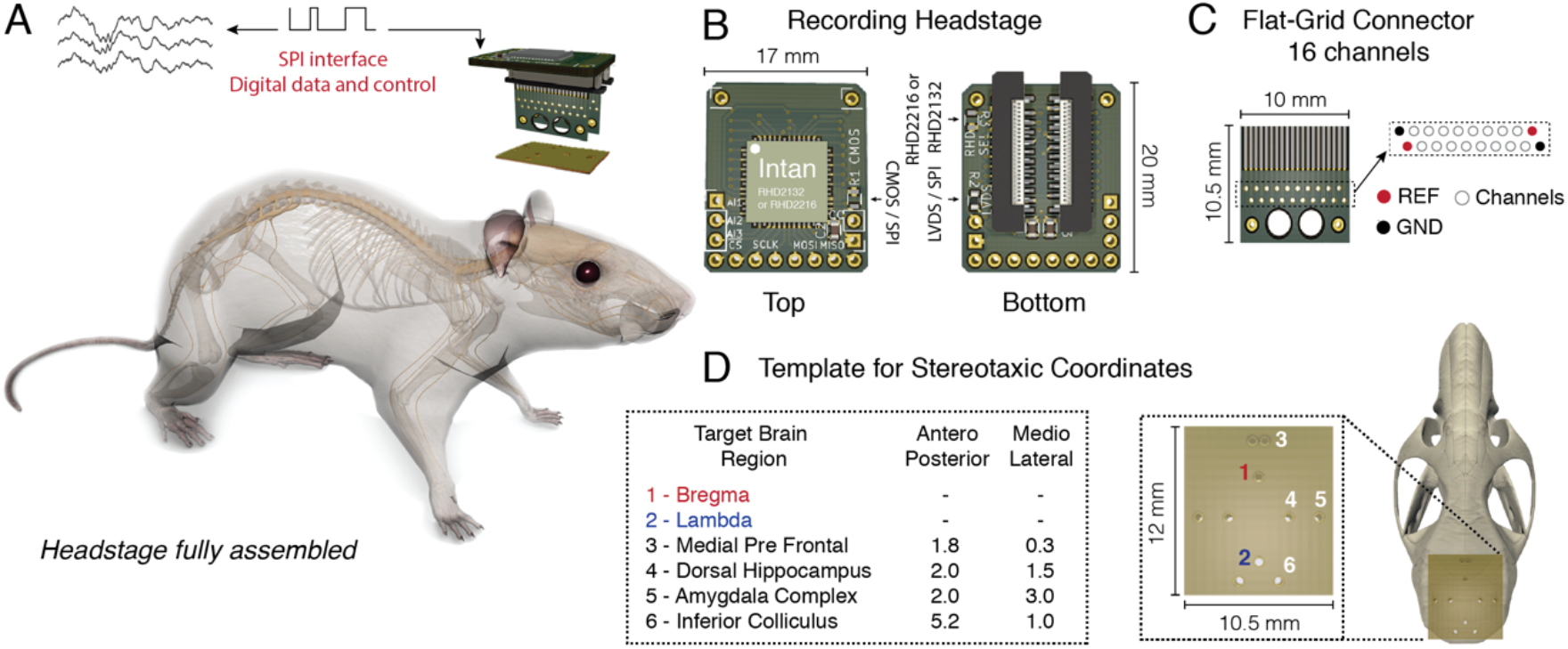
Headstage adapted from the Intan RHD headstage system with a flexible printed circuit (SMD/FPC) connector. A) Headstage fully assembled in real size. B) Top and bottom view of the headstage. 2132 or 2216 Intan chips can be used according to the circuit configuration. C) Flat-Grid connector board with sixteen channels, reference (REF) and ground (GND), designed to connect to the SMD/FPC connector. D) Fiber board punctured with small holes according to the chosen stereotaxic coordinates. A variable number of electrodes can be aligned and fixed from distinct dorsoventral configurations.

### Flat-Grid Connector

The modern technology of chronic electrodes most often involves Omnetics connectors that are implanted into the animal’s skull (Juavinett et al., 2019; Royer et al., 2010). After weeks of experimental procedures, the reuse of these connectors becomes almost impossible, since they are fixed with resistant cements. When removed, they are damaged either by the procedure or by the action of the solvents commonly used to dissolve the cemeteries.

To make chronic experiments more accessible we design a circuit board (10.5 mm x 10 mm x 0.4 mm) that works as a ‘rigid’ flat cable. Basically, at the top of this small PCB there is an interface to connect to the SMD/FPC connector, in the middle there is a grid of twenty contacts (through holes with 0.2 mm each separated by 0.5 mm; two grounds, two references and sixteen contacts for sixteen channels) where the electrodes are soldered, and at the bottom there are bigger holes to assist the cement fixation at the skull during implant surgery (Figure 1C).

### Template for Stereotaxic Coordinates

Most commercial electrodes usually have parallel distribution, that is, each shank is side by side and separated from each other by micrometers (Ulyanova et al., 2019). A similar parallel distribution has also been used by other authors who have sought to develop new and inexpensive alternatives for electrophysiological records (França et al., 2020).

In an attempt to develop a more versatile way that meets the multichannel recording of widely distributed neural networks, a fiber PCB substrate was punctured with small holes. Each hole is drilled with 400*μ*m precision and distributed as a function of an anteroposterior and mediolateral stereotaxic coordinate (Paxinos and Franklin, 2012), where a variable number of electrodes can be aligned and fixed from distinct dorsoventral configurations (e.g., Figure 1D).

### Wires

In this work we choose Tungsten microwires 99.5% S-Formvar with 50 μm internal diameter (California Fine Wire Co.) for the multi-wire arrays (Nicolelis, 2007) (Figure 2A). Tungsten wires are commonly used for electrophysiological records and are known for their high rigidity, durability, and high impedance (Hubel, 1957). Stainless steel with 0.127 mm internal diameter, Teflon-coated wires (Model 791400, A-M Systems Inc., Carlsborg, WA, USA) were used as reference and ground.

**Figure 2.**
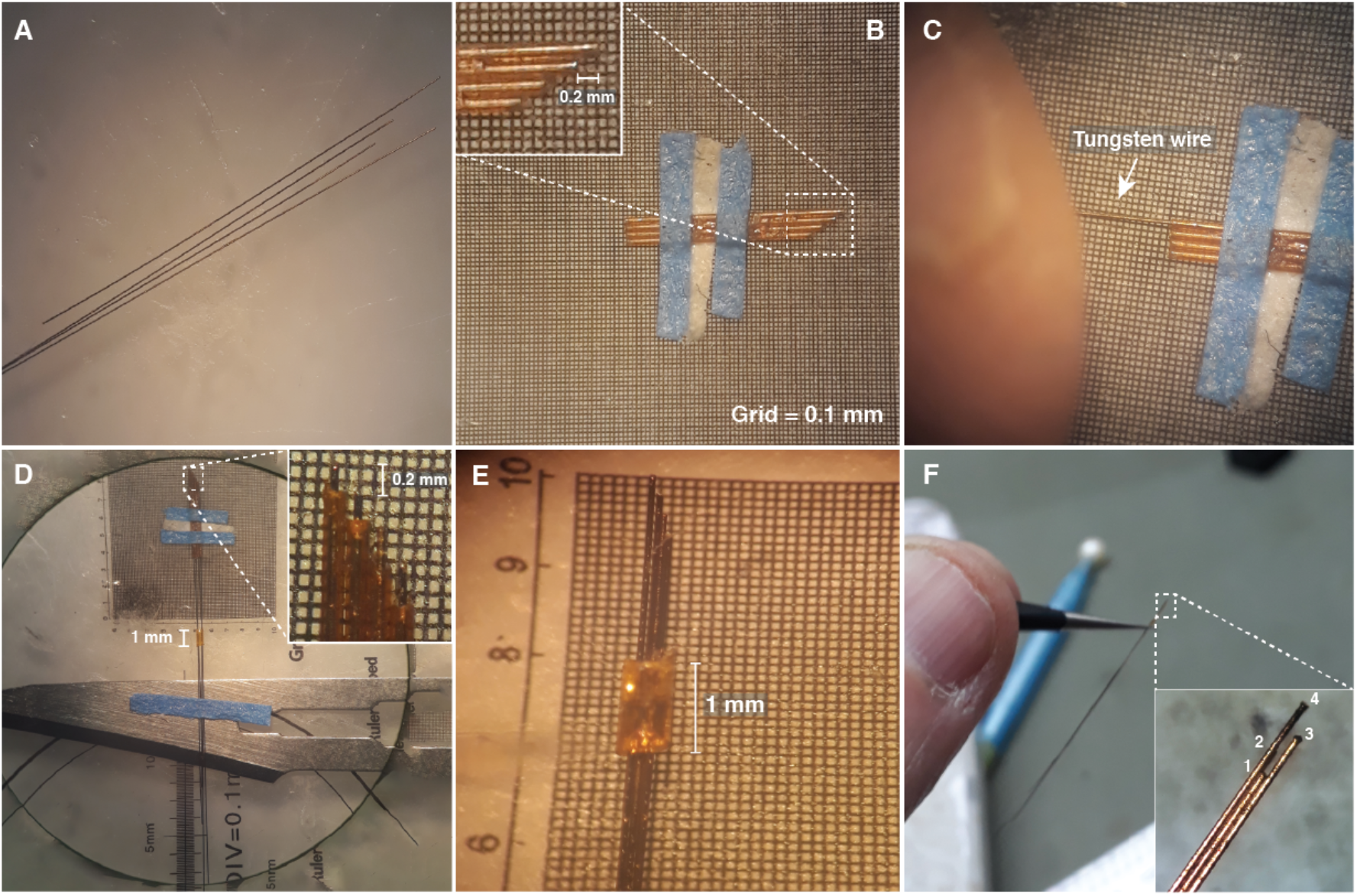
Designing electrode arrays. A) Tungsten wires 50 μm in diameter. B) Silica capillary tubes fixed side by side in a microscope micrometer calibration ruler, to form a matrix guide. C) Tungsten wire passing through the silica tube. D) Tungsten wires fixed with tips spaced by 200 μm. E) 1 mm silica tube used as a sliding sleeve to define the full length of the electrode array. F) Amplified electrode array showing the tip configuration.

### Assembly process

In order to record different brain regions and their respective neural subtracts, electrode arrays with several tungsten wires each could be built and attached to the template according to the previously established stereotaxic coordinates (e.g., Figure 1D). In turn, each wire tip from an electrode array could be spaced according to the experimental purpose (e.g., Figure 2B and 2E-F).

In what follows we describe one template type for stereotaxic coordinates that have been used in experiments in our laboratory. Each step of headstage manufacturing will be commented on below. More details can be found on supplementary material or GitHub (https://cutt.ly/PcrSqFY).

#### Step 1 - Electrode Array

1. Tungsten wires should be cut to different sizes. This will facilitate their identification later (Figure 2A)
2. For the sizing of the electrode arrays, fused silica capillary tubes 75 μm internal diameter and 155 μm external diameter (Polymicro Technologies - https://cutt.ly/6x6yXlM) were fixed side by side in a microscope micrometer calibration ruler, to form a matrix guide. This matrix was used for the correct wire alignment and to ensure a specific distance between each wire tip. This distance was in accordance with the dorsal-ventral stereotaxic coordinates of the chosen brain substrates (Figure 2B).
3. Each tungsten wire should pass in isolation in each silica tube. Do not use tweezers or any tool that could compromise the wire insulation (Figure 2C).
4. A fused silica tube with 1 mm length, 200 μm internal diameter, and 350 μm external diameter should hold all wires on the rear surface. The wires, in turn, must be fixed to a removable surface (Figure 2D).
5. The 1 mm silica tube should be used as a sliding sleeve, that positioned on the rule, will define the full length of the electrode array. After the desired positioning, a small drop of super glue can be used to hold the wires next to the tube (Figure 2E).

#### Step 2 - Fixing the electrode arrays to the template

1. The fiber board (thickness: 1 mm) needs to be punctured with small holes (PCB drill bits - 0.4 mm), according to the stereotaxic coordinates chosen for the electrode array implant (Figure 3A). A support base to hold the fiber board was designed on a 3D printer, and this can in turn can be fixed on the stereotaxic frame (Figure 3A and 3B; Supplementary Video 1; https://cutt.ly/ecAM15O).
2. On the stereotaxic frame, each electrode array should be positioned in the respective coordinate previously punctured on the board (Figure 3B, 3C and 3D; Supplementary Video 2). After the desired positioning, a small drop of super glue can be used to hold the electrode arrays next to the board.

**Figure 3.**
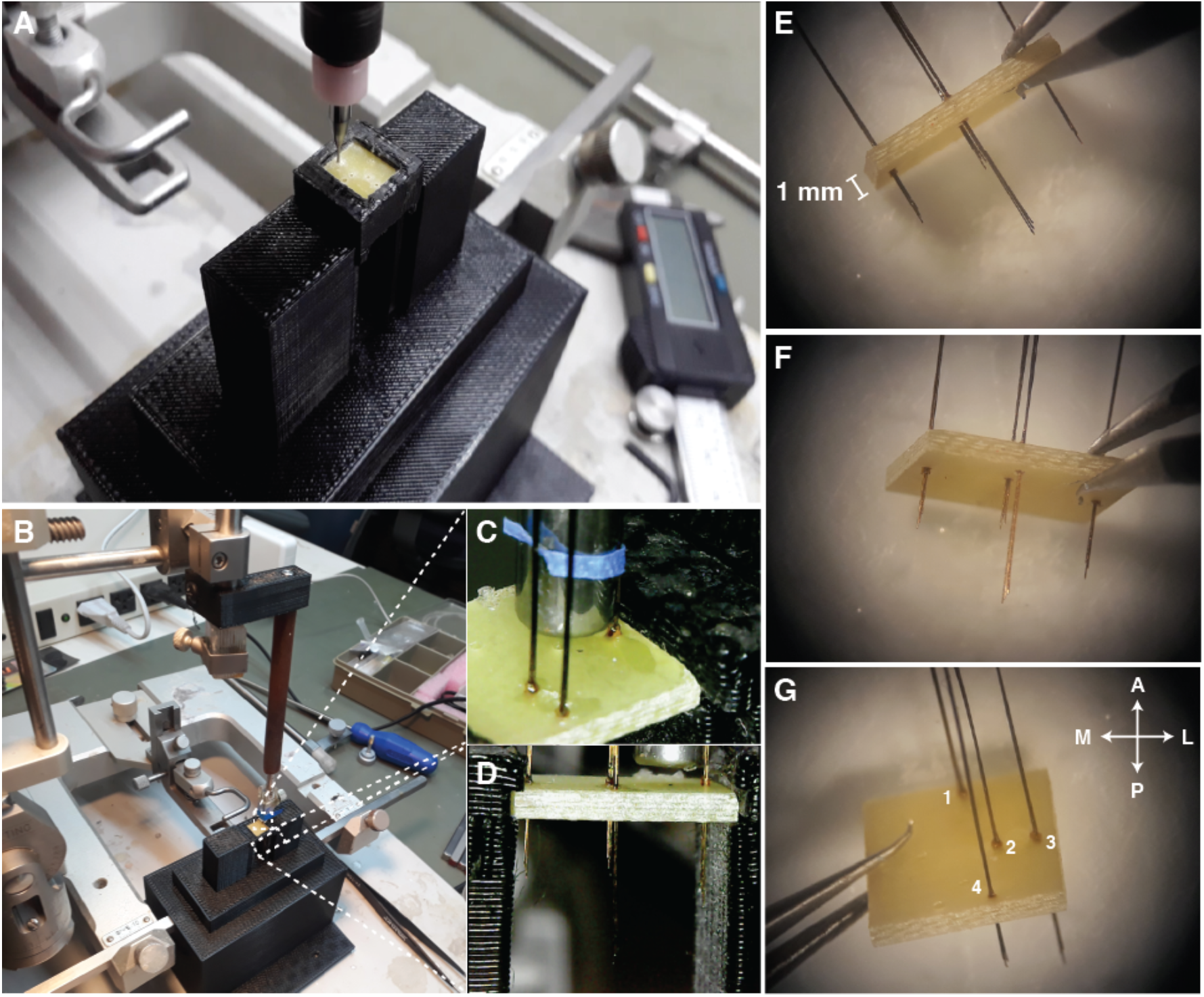
Fixing the electrodes to the template. A) Support base designed to hold the fiber board while being drilled by a drill. B) Support base attached to the stereotaxic frame. C) Each electrode array must be carefully positioned at each coordinate and glued with a small drop of super glue. D) Lateral view of the positioned electrode arrays. E-G) Template prepared with all electrode arrays. Antero <=> Posterior (A <--> P), Medial <=> Lateral (M <=> L) views.

#### Step 3 - Soldering the electrode arrays to the Flat-Grid Connector

1. A small support base to hold both the flat-grid connector and the fiber board, with pre-prepared electrode arrays, was designed using a 3D printer. This support ensures that tungsten wires will not bend during any manipulation. In addition, the holder facilitates both soldering and fixing wires over the grid (Figure 4A; https://cutt.ly/ecAM15O).
2. After connecting the flat-grid connector and the fiber board in the holder a small drop of super glue can be used to hold both (Figure 4B).
3. With a tweezer, make the insertion of each tungsten wire through the grid contacts. Be careful with the positioning of each electrode, in the correct channel contact. Remember that each electrode has a different length and this can be used as a position marker. (Figure 4C).
4. In the same way, as in item 3, put the wires on the reference and ground positions on the left and right sides of the grid. Note that the positioning of the ground and reference can be performed alternately (ground on the left and reference on the right or vice versa) (Figure 1C and 4D).
5. With a sharp blade remove the insulation of each wire next to the contact on the grid (Figure 4E).
6. Use a pointed solder iron with a small amount of weld to cover the contact between the tungsten wire and the grid (Figure 1F). Be aware that Tungsten is a refractory metal that needs special conditions to be soldered. Thus, we recommend using silver paint (SPI Supplies^Ⓡ^) to ensure a short-circuit between the tungsten wire and the grid contact (Figure 1G). In our experience, both methods should be used because the weld does not only fix the wire but the heat ensures the melting of the insulation.
7. Remove the parts from the support base and do the same procedures described in items 5 and 6 on the back of the flat-grid connector. Cut the wires close to the grid (Figure 4H).
8. Cover all grid contacts, on both sides, with a thermosetting polymer (Epoxy resin, Devcon^Ⓡ^) (Figure 4I).
9. Optionally, the fiber board corners can be removed with a rotatory tool (e.g., mini cordless electric drill with a sanding band) to fit better on the animal skull (Figure 4J).

**Figure 4.**
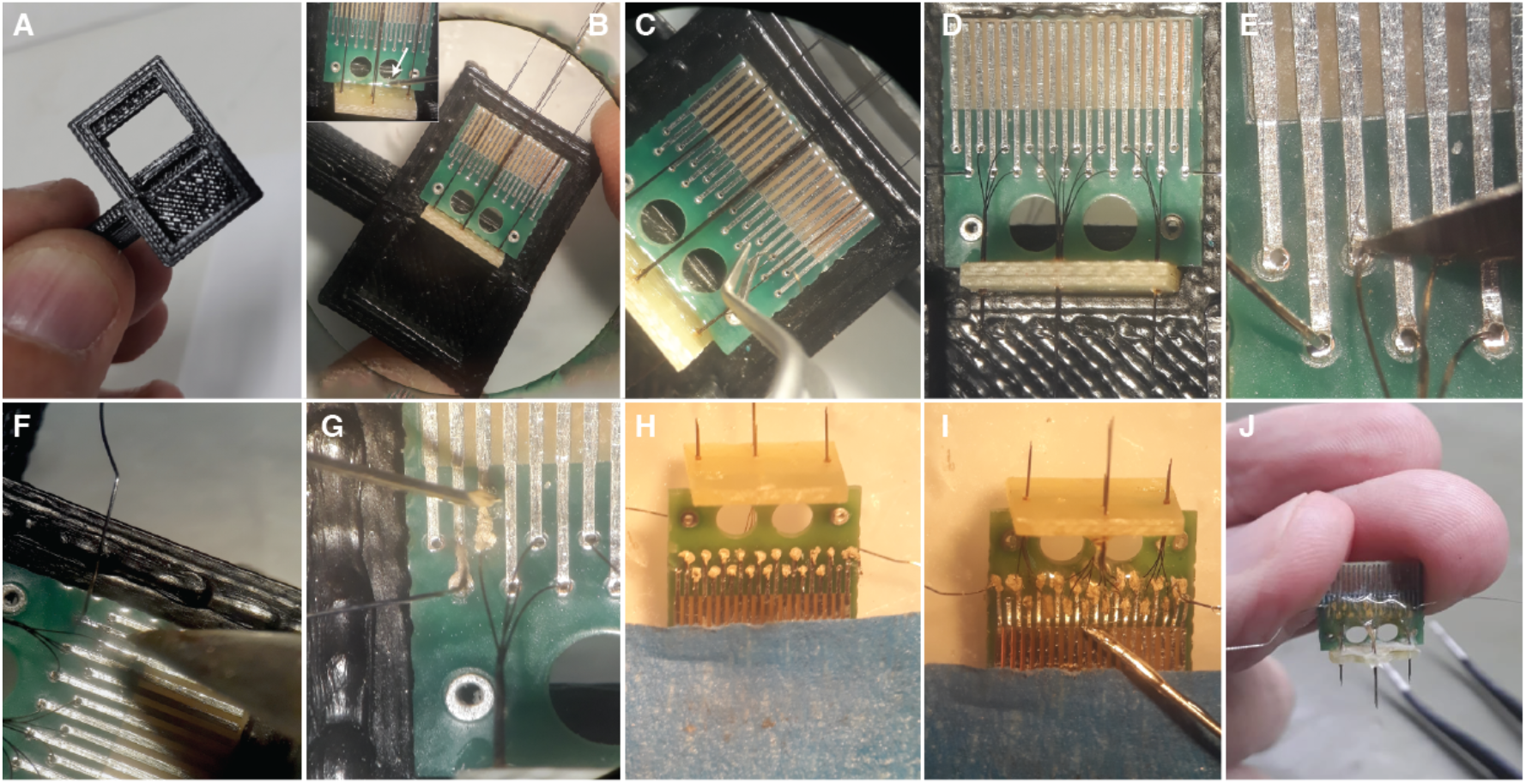
Preparing the Flat-Grid Connector. A-B) Support base to hold both the flat-grid connector and the fiber board. C-D) Each wire needs to be inserted through the grid contacts. E) The insulation of each wire needs to be removed. F) Small amount of weld covering the contact between the wire and the grid. G) Silver paint applied to short-circuit the contact. H) Electrode arrays and Flat-Grid Connector at the posterior and bottom view. Wires cut close to the grid. I) Grid contacts covered by a thermosetting polymer. J) Finished set with board corners removed.

#### Step 4 - Electroplating

In order to reduce the impedance imposed by the small tungsten wire diameter, each tip was electroplated with gold (9V, 0.075 mA; Direct Current over 60s) in a handmade system built for this purpose (https://cutt.ly/QcrUCvL). The impedance was measured in ACSF (artificial cerebrospinal fluid) solution at 25 °C [NaCl (127 mM), KCl (2 mM), CaCl_2_ (2 mM), MgSO_4_ (2 mM), NaHCO_3_ (26 mM), KH_2_PO_4_ (1.2 mM), HEPES (13 mM); pH 7.4], by the Intan RHD2132 via Open Ephys acquisition system (Siegle et al., 2017) (Table 1).

**Table 1:** Impedance measured by INTAN RHD2132

| Ch | 1 | 2 | 3 | 4 | 5 | 6 | 7 | 8 | 9 | 10 | 11 | 12 | 13 | 14 | 15 | 16 |
| --- | --- | --- | --- | --- | --- | --- | --- | --- | --- | --- | --- | --- | --- | --- | --- | --- |
| Z | 6.6 | 5.5 | 3.8 | 5.6 | 4.5 | 30.2 | 6.8 | 3.2 | 92.0 | 63.5 | 5.1 | 11.7 | 28.0 | 24.5 | 9.7 | 7.7 |

Ch - channels; Z - impedance in $K\Omega$

### Ethical statement

The data and experimental procedures shown in this paper are from two male mice (C57BL/6 strain) recorded during two different experiments carried out in our laboratory.

All procedures were approved by the Institutional Animal Care and Use Committee at the Universidade Federal de Minas Gerais (CEUA-UFMG: 198/2019), conducted in accordance with Conselho Nacional de Controle de Experimentação Animal (CONCEA) guidelines defined by Arouca Act 11.794 under Brazilian federal law. CEUA directives comply with National Institutes of Health (NIH) guidelines for the care and use of animals in research.

### Surgical procedure

The mice were anesthetized with Isoflurane in Oxygen at concentrations of 2–4% for induction and 0.5–2.0 % for maintenance. After the absence of reflexes and signs of pain, the head surface was shaved and then the animal was positioned in a stereotaxic frame (Stoelting, Wood Dale, IL, United States). The constant flow of anesthetic was offered through a 3D printed mask designed in our laboratory. After asepsis with alcohol (70%, topical) and povidineiodine solution (7.5%, topical), local anesthesia with lidocaine clorohydrate-epinephrine [1% (wt/vol), 7 mg/kg] was applied and the scalp was removed to expose the skull.

Small holes at the top of the skull were made with a 0.5 mm drill according to stereotaxic coordinates previously defined in the template board (Figure 1D), in turn, the dorso-ventral coordinates were defined according to the length of the electrode arrays. Each array was designed with four electrodes, with tips spaced by 200 μm (Figure 2).

In mouse 1, four electrode arrays were distributed as follows: Medial Pre-Frontal Cortex (mPFC), 3.0 mm; Amygdala Complex (AMY), 4.8 mm; Dorsal Hippocampus (dHPC), 2.0 mm; Inferior Colliculus (IC), 2.2 mm (Figure 1D and Figure 5). In mouse 2, one electrode array was positioned only in the Inferior Colliculus (IC), 2.2 mm. At the end of the procedure, the implant was fixed to the skull with dental acrylic (Figure 5).

**Figure 5.**
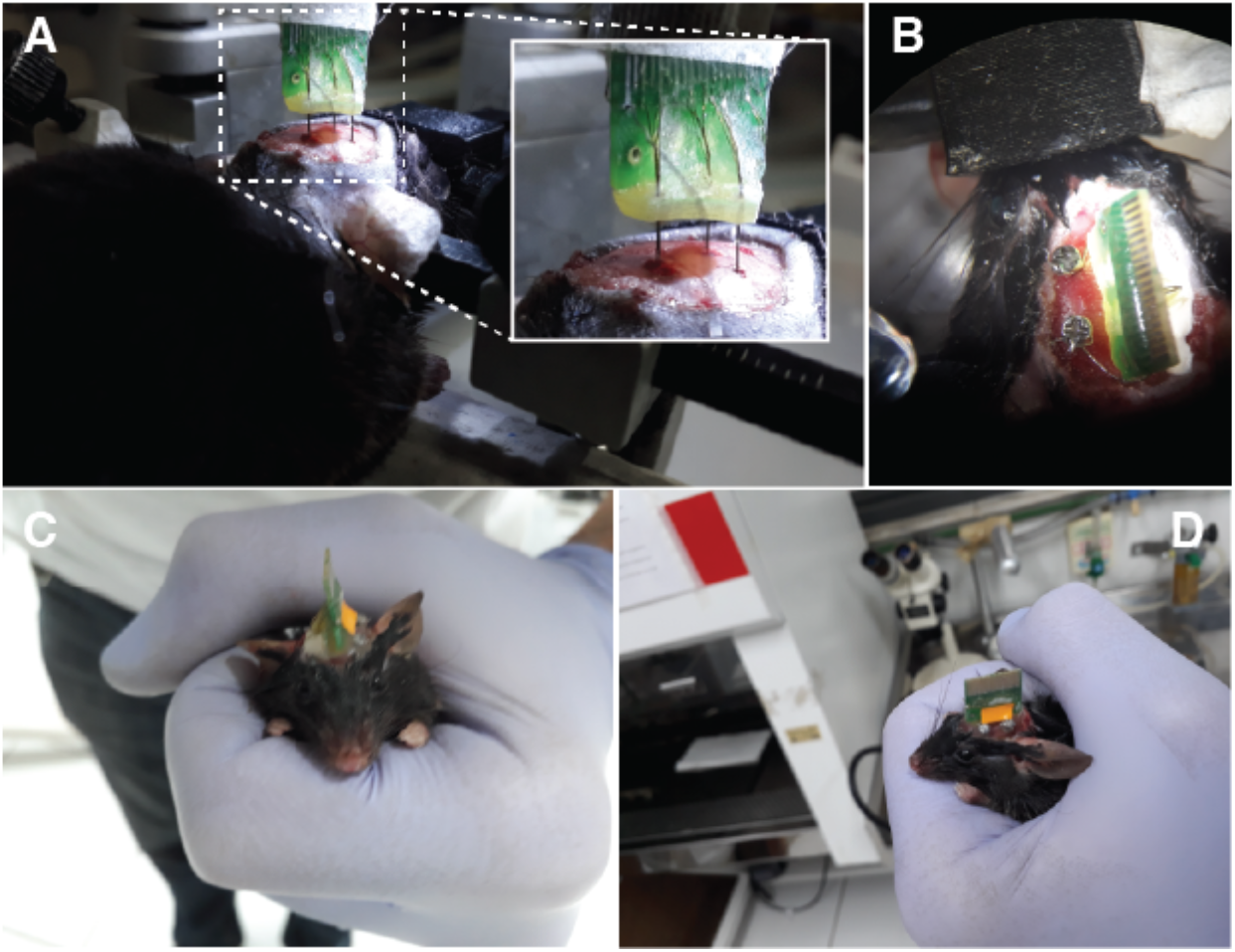
Surgical procedure. A) Electrode arrays being positioned in the animal brain. B) Top view. Fixing screws used as a reference and ground. C-D) Animal after surgery. Anterior and lateral views, respectively.

Following the surgery, animals received an intramuscular injection of penicillin-G benzathine, a broad-spectrum antibiotic cocktail (Pentabiotic^®^, 56.7 mg/kg in a 0.1 mL volume); a subcutaneous injection of anti-inflammatory analgesic (Banamine^®^; 0.5 mg/kg flunixin meglumine in a volume of 0.3 mL) and a subcutaneous injection of opioid analgesic (Tramadol^®^; 20 mg/kg s.c. in a volume of 0.1 mL) and were allowed a recovery period of seven days.

### Electrophysiological Record and Behavioral Protocol

The raw electrophysiological signals were obtained from the Intan RHD2132 via Open Ephys acquisition system. All records were sampled at 30 kHz and stored as continuous data files (*. continuous) (Siegle et al., 2017).

The animals used in the experiment were placed in a square open-field arena (30 cm × 30 cm × 30 cm) where an auditory stimuli protocol was performed. The protocol consisted of five auditory stimuli that evoked an entrainment by applying a long-lasting amplitude-modulated tone. This auditory stimulation is currently used in our laboratory for learning and memory tasks (Lockmann et al., 2017; Simões et al., 2020).

A 12-bit digital-to-analog converter at Arduino Due was used to generate the auditory stimuli (Amaral-Júnior et al., 2019; Picton et al., 2003). The algorithm behind the tone generation is openly available (https://cutt.ly/FzHgv7H). The stimuli consisted of a 10 kHz pure tone applied for 30 s and modulated at an amplitude of 53.71 Hz sine wave (100 % modulation depth) and set to 85 dB SPL (Brüel & Kjaer type 2238 sound level meter).

The timestamps locked to the peaks and valleys of the modulated tone were recorded through the Open Ephys digital input port. Via a linear interpolation, these time values were used to obtain an instantaneous phase time series, which in turn was used to reconstruct the signal. This allowed the time-frequency analysis to remain engaged to the auditory stimulation (Amaral-Júnior et al., 2019).

Finally, to track the animal’s movement, the whole experiment was filmed (640×480 resolution; 30 frames per second) by a camera (Logitech^®^, C270 Hd 720p) set at the top of the arena.

### Data analysis

The data were analysed off-line using custom-made codes and standard MATLAB functions (MATLAB R2020a Signal Processing Toolbox and EEGlab toolbox - https://sccn.ucsd.edu/eeglab/index.php). For the analysis of local field potentials (LFP), the signal was decimated to 3 kHz. Time-frequency power and the power spectrum density of the evoked potentials were analyzed by using the standard spectrogram (non-overlapping, 16384-point, Hamming window) and pwelch (non-overlapping, 4096-point, Hamming window) Matlab functions, respectively. For the analysis of neuronal units, the signal was processed in an unsupervised algorithm for spike detection and sorting (Chaure et al., 2018).

The open-source software Bonsai (Lopes et al., 2015) was used to track the animal over the arena. From the x and y coordinates of each video frame, we created a tracking map and calculated the displacement and the total distance traveled.

## RESULTS

### Local Field Potentials

Long-lasting modulated tones can evoke a stable oscillatory activity at different levels of the auditory system (Picton et al., 2003). More specifically, the IC can be entrained by a programmed frequency modulation (Lockmann et al., 2017; Simões et al., 2020).

As expected, over five stimulus presentations, the oscillatory activity in the ventral IC showed a distinct power spectral signature at the same modulated frequency (53.7 Hz) (Figure 6C - Top and Supplementary Figure 2). Moreover, the mean power value at the programmed frequency modulation was quite representative in the frequency spectrum (Figure 6D - Top). It is worth mentioning that, according to the IC tonotopic organization, central and ventral IC perfectly responds to high-frequency stimulation (10 kHz) (Clopton and Winfield, 1973; Malmierca et al., 2008).

**Figure 6.**
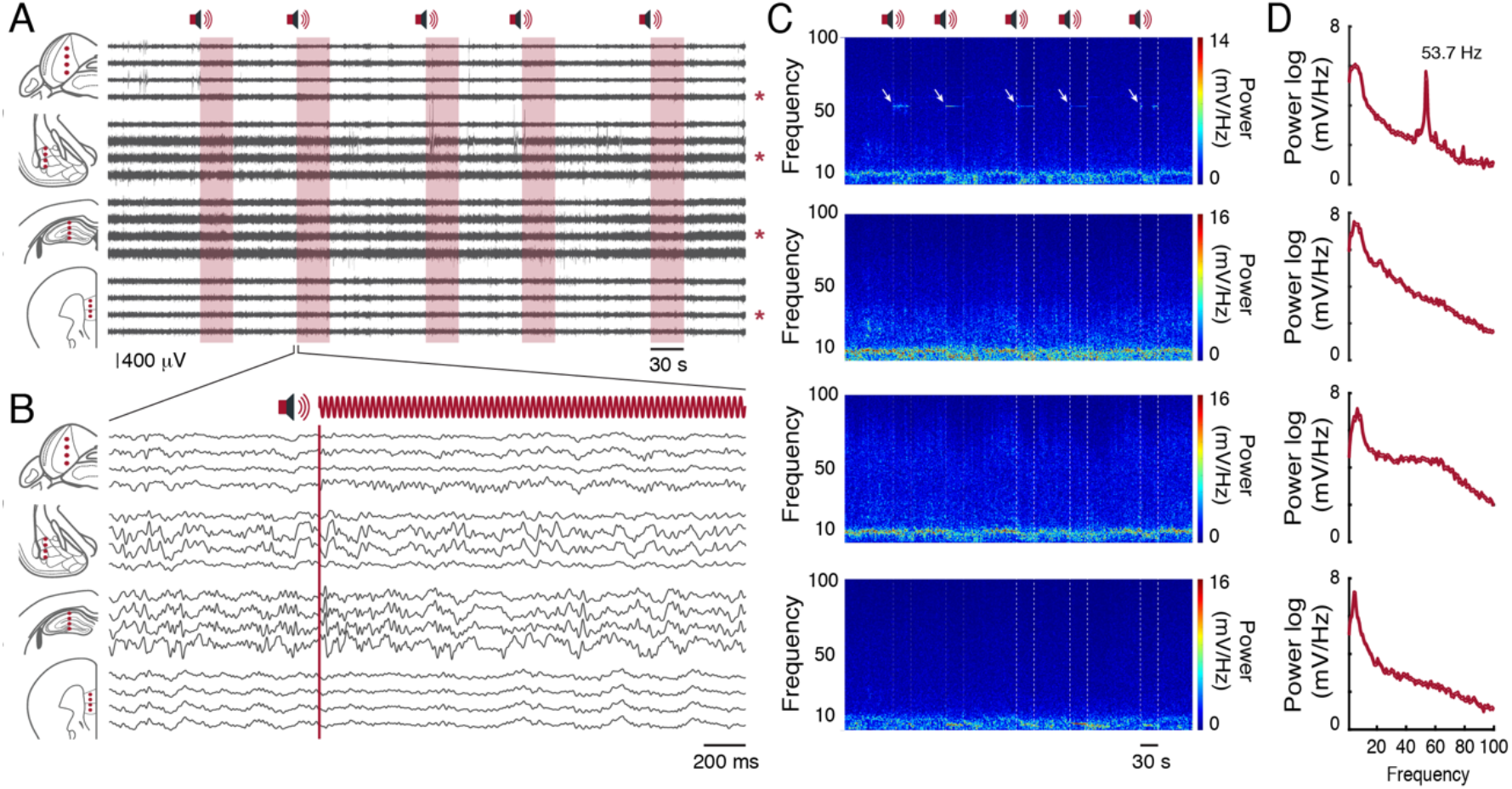
Local Field Potentials. A) Raw record of sixteen channels on each brain substrate. In red we have the auditory stimulus presentation window. Pictorial representations of channels positioned in each brain region are represented on the left. Each red asterisk on the right represents the channel chosen for the time-frequency analysis. B) Two-second time window of the second auditory stimulus presentation. Filtered records between 1 and 100 Hz. C) Channel time-frequency power spectrogram signaled by red asterisks in A. From top to bottom: IC, AMY, dHPC, mPFC. D) Channel mean power spectral density of the evoked potentials signaled by red asterisks in A. From top to bottom: IC, AMY, dHPC, mPFC.

As described above, Mouse 1 had four electrode arrays distributed over the IC, AMY, dHPC, and mPFC. Each electrode array had four tungsten wires in four dorsal-ventral coordinates (Figure 2F). According to the records, each brain region showed a distinct oscillatory pattern, without visible crosstalk contamination between channels (Figure 6B and 6C). While the largest amplitude theta oscillations were observed at the dHPC (Buzsáki, 2002), theta and other rhythms can also be observed at both substrates with different features (Cohen et al., 2021; Soltani Zangbar et al., 2020).

### Single Units

We identified four cell types in the ventral IC that substantially changed the activity to the auditory stimuli (Figure 7). Interestingly, two cells exhibit a continuous firing, but while one increases the firing rate in front of the stimulus (Figure 7C and 7D - red cell), the other one keeps the firing rate constant, changing its timing according to the modulation frequency period (Figure 7C and 7E - blue cell). The other two cells respond only in front of the stimulus (Figure 7C and 7F - green cell; Figure 7C and 7G - orange cell).

**Figure 7.**
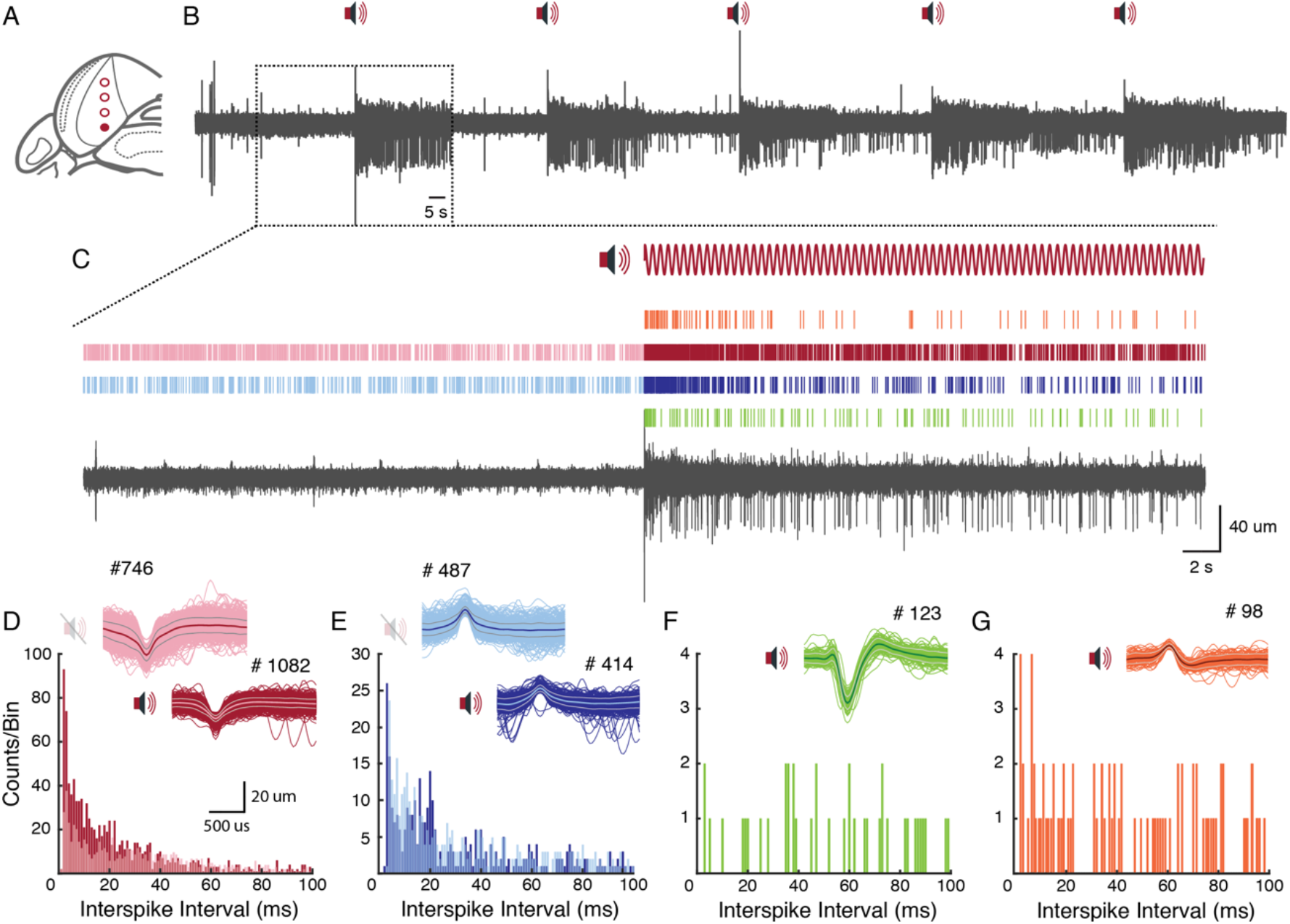
Single units sorting. A) Pictorial representation of channels positioned in the IC. The dot filled in red represents the channel chosen for the analysis. B) Full record filtered between 300 and 3000 Hz with the respective auditory stimuli presentations. C) Sixty-second time window, before and during the auditory stimulus presentation. Above the record is represented four types of cells and their respective firing rate over time. D-E) Interspike-interval histogram of each cell type. Above the histograms, the respective waveform and the total firing rate are represented.

Prominent single units can also be qualitatively observed along the sixteen recorded channels (Supplementary Figure 1).

### Exploratory behavior

The exploratory behavior was analyzed in order to identify any impairment of the animal’s locomotion, in face of surgery and/or the new headstage design. However, in both animals, the accumulated distance and the displacement over time showed a linear (Figure 8B and 8E) and constant distribution (Figure 8C and 8F), with distances covered by more than 10 meters at the end of the protocol. In general, during the experimental protocol, the animals did not show signs associated with pain, discomfort and/or distress (Supplementary Video 3).

**Figure 8.**
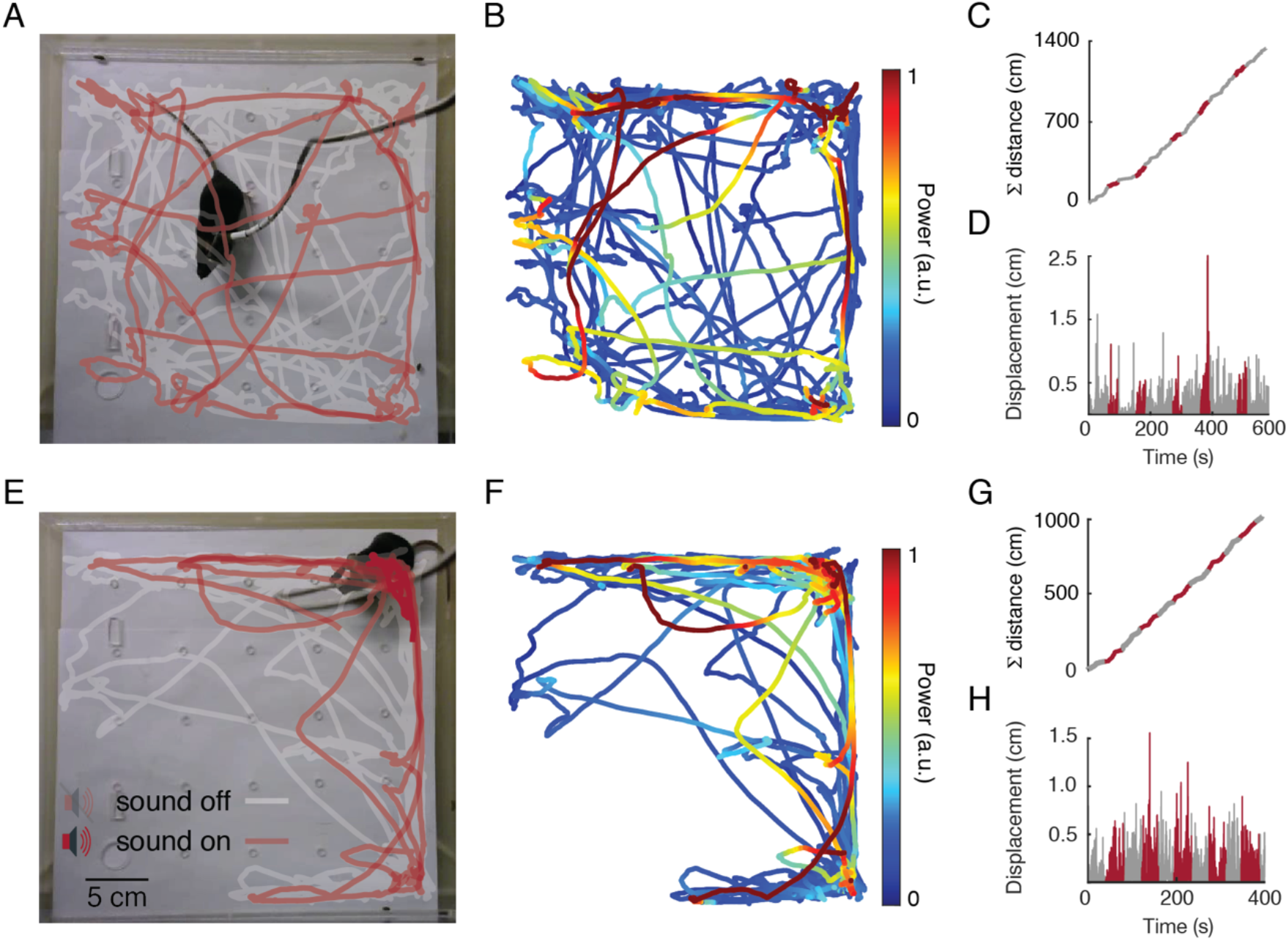
Exploratory behavior analysis. Mouse 1 and Mouse 2 are represented at the top and bottom of the figure, respectively. A) and E) Tracking maps. B) and F) Ventral IC power spectral at the same modulated frequency band (53.7 ± 0.3 Hz) on the tracking map. C) and G) Accumulated distance over time. D) and H) Displacement over time.

## DISCUSSION

Over the past few years, open-source initiatives have encouraged the development of unique investigation methods that are accessible to research groups. Impressively, some brilliant scientific teams have developed and shared cutting-edge methodologies (Lopes et al., 2015; Mathis et al., 2018) and equipment (Siegle et al., 2017) free of charge or made available at nominal cost. It is worth mentioning that not relying on ordinary equipment, that does not provide any knowledge of its internal workings, is an important step to ensure reproducibility and accuracy with a high degree of agreement.

Based on this democratic initiative of accessible and equitable science, which involves developing more affordable materials and methods, we designed an adaptation to the RHD headstage system (Intan technologies), associated with a flat connector (SMD/FPC) to replace the traditional Omnetics connector. It is unquestionably that the Omnetics connectors have a high technological level and are widely used by the scientific community in several applications. However, these may lead to prohibitive costs for underfunded research groups (especially in developing countries) and are a limiting factor when designing experimental protocols. On the other hand, the flat connector has affordable costs, is easily recycled, and can be also easily adapted to the experiments.

For proper connection with the new connector, we designed a small circuit board with a through holes grid, where the electrodes are soldered. Beside that, a complete structure for the placement of the electrode arrays was designed to adapt to the specific needs of different projects.

According to our results, the experimental animals easily adapted to the headstage design, without discomfort or impairments not interfering in its mobility and allowing the task execution. Besides, the extracellular records had great quality with detectable evoked responses to an auditory task both on local field and unit activities.

In Neuroscience, we believe that versatile and flexible tools allow the study of biological processes that are not observable with ordinary equipment. Moreover, the development of customized equipment can enable local scientific development, as it inspires scientific endeavor among students, teachers, and supervisors alike. Our project is available through open-source initiatives (https://cutt.ly/PcrSqFY) and may be modified by collaborators when adding new features to both developed segments.

## ACKNOWLEDGMENTS

We would like to thank Cristiano Soares Simões, Marina Pádua Reis and John Warrener for the constructive criticism that led to improvements in the manuscript and Davi Barreto Mourão and Frederico José Barreto Mourão for all inspiration.

## AUTHOR CONTRIBUTIONS STATEMENT

Projected conceived and designed by F.A.G.M., L.O.G., P.A.A.J. and M.F.D.M.; F.A.G.M., L.O.G., P.A.A.J., V.R.C. and M.F.D.M. designed boards and connectors; F.A.G.M., performed surgery and electrophysiological records; F.A.G.M. and L.O.G., performed behavioral tests; F.A.G.M., performed electrophysiological and behavioral analysis; F.A.G.M. prepared figures; E.M.A.M.M. supervised and contributed to refinements of the system; F.A.G.M. drafted the manuscript; All authors revised and approved the final version of manuscript.

## REFERENCES

Amaral-Júnior, P.A., Mourão, F.A.G., Amancio, M.C.L., Pinto, H.P.P., Carvalho, V.R., Guarnieri, L. de O., Magalhães, H.A., Moraes, M.F.D., 2019. A Custom Microcontrolled and Wireless-Operated Chamber for Auditory Fear Conditioning. Front. Neurosci. 13, 1193.

Buzsáki, G., 2002. Theta oscillations in the hippocampus. Neuron 33, 325–340.

Buzsáki, G., Stark, E., Berényi, A., Khodagholy, D., Kipke, D.R., Yoon, E., Wise, K.D., 2015. Tools for probing local circuits: high-density silicon probes combined with optogenetics. Neuron 86, 92–105.

Chaure, F.J., Rey, H.G., Quian Quiroga, R., 2018. A novel and fully automatic spike-sorting implementation with variable number of features. J. Neurophysiol. 120, 1859–1871.

Chung, J.E., Joo, H.R., Fan, J.L., Liu, D.F., Barnett, A.H., Chen, S., Geaghan-Breiner, C., Karlsson, M.P., Karlsson, M., Lee, K.Y., Liang, H., Magland, J.F., Pebbles, J.A., Tooker, A.C., Greengard, L.F., Tolosa, V.M., Frank, L.M., 2019. High-Density, Long-Lasting, and Multi-region Electrophysiological Recordings Using Polymer Electrode Arrays. Neuron 101, 21–31.

Clopton, B.M., Winfield, J.A., 1973. Tonotopic organization in the inferior colliculus of the rat. Brain Res. 56, 355–358.

Cohen, M.X., Englitz, B., França, A.S.C., 2021. Large- and multi-scale networks in the rodent brain during novelty exploration. eNeuro. 0494-20.2021

Dowben, R.M., Rose, J.E., 1953. A metal-filled microelectrode. Science 118, 22–24.

França, A.S.C., van Hulten, J.A., Cohen, M.X., 2020. Low-cost and versatile electrodes for extracellular chronic recordings in rodents. Heliyon 6, e04867.

Green, J.D., 1958. A Simple Microelectrode for recording from the Central Nervous System. Nature. https://doi.org/10.1038/182962a0

Hallal, P.C., 2021. SOS Brazil: science under attack. Lancet 397, 373–374.

Harrison, R.R., 2007. A versatile integrated circuit for the acquisition of biopotentials. Custom Integrated Circuits Conference, CICC’07. IEEE 2008, 115–122.

Hong, G., Lieber, C.M., 2019. Novel electrode technologies for neural recordings. Nat. Rev. Neurosci. 20, 330–345.

Hubel, D.H., 1957. Tungsten Microelectrode for Recording from Single Units. Science. https://doi.org/10.1126/science.125.3247.549

Jones, K.E., Campbell, P.K., Normann, R.A., 1992. A glass/silicon composite intracortical electrode array. Ann. Biomed. Eng. 20, 423–437.

Juavinett, A.L., Bekheet, G., Churchland, A.K., 2019. Chronically implanted Neuropixels probes enable high-yield recordings in freely moving mice. Elife 8. https://doi.org/10.7554/eLife.47188

Jun, J.J., Steinmetz, N.A., Siegle, J.H., Denman, D.J., Bauza, M., Barbarits, B., Lee, A.K., Anastassiou, C.A., Andrei, A., Aydın, Ç., Barbic, M., Blanche, T.J., Bonin, V., Couto, J., Dutta, B., Gratiy, S.L., Gutnisky, D.A., Häusser, M., Karsh, B., Ledochowitsch, P., Lopez, C.M., Mitelut, C., Musa, S., Okun, M., Pachitariu, M., Putzeys, J., Rich, P.D., Rossant, C., Sun, W.-L., Svoboda, K., Carandini, M., Harris, K.D., Koch, C., O’Keefe, J., Harris, T.D., 2017. Fully integrated silicon probes for high-density recording of neural activity. Nature 551, 232–236.

Lockmann, A.L.V., Mourão, F.A.G., Moraes, M.F.D., 2017. Auditory fear conditioning modifies steady-state evoked potentials in the rat inferior colliculus. J. Neurophysiol. 118, 1012–1020.

Lopes, G., Bonacchi, N., Frazão, J., Neto, J.P., Atallah, B.V., Soares, S., Moreira, L., Matias, S., Itskov, P.M., Correia, P.A., Medina, R.E., Calcaterra, L., Dreosti, E., Paton, J.J., Kampff, A.R., 2015. Bonsai: an event-based framework for processing and controlling data streams. Front. Neuroinform. 9, 7.

Malmierca, M.S., Izquierdo, M.A., Cristaudo, S., Hernández, O., Pérez-González, D., Covey, E., Oliver, D.L., 2008. A discontinuous tonotopic organization in the inferior colliculus of the rat. J. Neurosci. 28, 4767–4776.

Mathis, A., Mamidanna, P., Cury, K.M., Abe, T., Murthy, V.N., Mathis, M.W., Bethge, M., 2018. DeepLabCut: markerless pose estimation of user-defined body parts with deep learning. Nat. Neurosci. 21, 1281–1289.

McNaughton, B.L., O’Keefe, J., Barnes, C.A., 1983. The stereotrode: a new technique for simultaneous isolation of several single units in the central nervous system from multiple unit records. J. Neurosci. Methods 8, 391–397.

Nicolelis, M.A.L., 2007. Methods for Neural Ensemble Recordings, Second Edition. CRC Press.

Paxinos, G., Franklin, K.B.J., 2012. Paxinos and Franklin’s the Mouse Brain in Stereotaxic Coordinates. Academic Press.

Picton, T.W., John, M.S., Dimitrijevic, A., Purcell, D., 2003. Human auditory steady-state responses. Int. J. Audiol. 42, 177–219.

Polo-Castillo, L.E., Villavicencio, M., Ramírez-Lugo, L., Illescas-Huerta, E., Moreno, M.G., Ruiz-Huerta, L., Gutierrez, R., Sotres-Bayon, F., Caballero-Ruiz, A., 2019. Reimplantable Microdrive for Long-Term Chronic Extracellular Recordings in Freely Moving Rats. Front. Neurosci. 13, 128.

Rousche, P.J., Normann, R.A., 1998. Chronic recording capability of the Utah Intracortical Electrode Array in cat sensory cortex. J. Neurosci. Methods 82, 1–15.

Royer, S., Zemelman, B.V., Barbic, M., Losonczy, A., Buzsáki, G., Magee, J.C., 2010. Multi-array silicon probes with integrated optical fibers: light-assisted perturbation and recording of local neural circuits in the behaving animal. Eur. J. Neurosci. 31, 2279–2291.

Siegle, J.H., López, A.C., Patel, Y.A., Abramov, K., Ohayon, S., Voigts, J., 2017. Open Ephys: an open-source, plugin-based platform for multichannel electrophysiology. J. Neural Eng. 14, 045003.

Simões, C.S., Mourão, F.A.G., Guarnieri, L.O., Passos, M.C., Moraes, M.F., 2020. Amygdala inhibition impairs fear conditioning but increases the stimulus-driven activity in the inferior colliculus. Neurosci. Lett. 738, 135311.

Soltani Zangbar, H., Ghadiri, T., Seyedi Vafaee, M., Ebrahimi Kalan, A., Fallahi, S., Ghorbani, M., Shahabi, P., 2020. Theta Oscillations Through Hippocampal/Prefrontal Pathway: Importance in Cognitive Performances. Brain Connect. 10, 157–169.

Szostak, K.M., Grand, L., Constandinou, T.G., 2017. Neural Interfaces for Intracortical Recording: Requirements, Fabrication Methods, and Characteristics. Front. Neurosci. 11, 665.

Ulyanova, A.V., Cottone, C., Adam, C.D., Gagnon, K.G., Cullen, D.K., Holtzman, T., Jamieson, B.G., Koch, P.F., Chen, H.I., Johnson, V.E., Wolf, J.A., 2019. Multichannel Silicon Probes for Awake Hippocampal Recordings in Large Animals. Front. Neurosci. 13, 397.

Voigts, J., Siegle, J.H., Pritchett, D.L., Moore, C.I., 2013. The flexDrive: an ultra-light implant for optical control and highly parallel chronic recording of neuronal ensembles in freely moving mice. Front. Syst. Neurosci. 7, 8.

