## Supplementary for "A fully adapted headstage for electrophysiological experiments with custom and scalable electrode arrays to record widely distributed brain regions": Supplementary_Figures.pdf

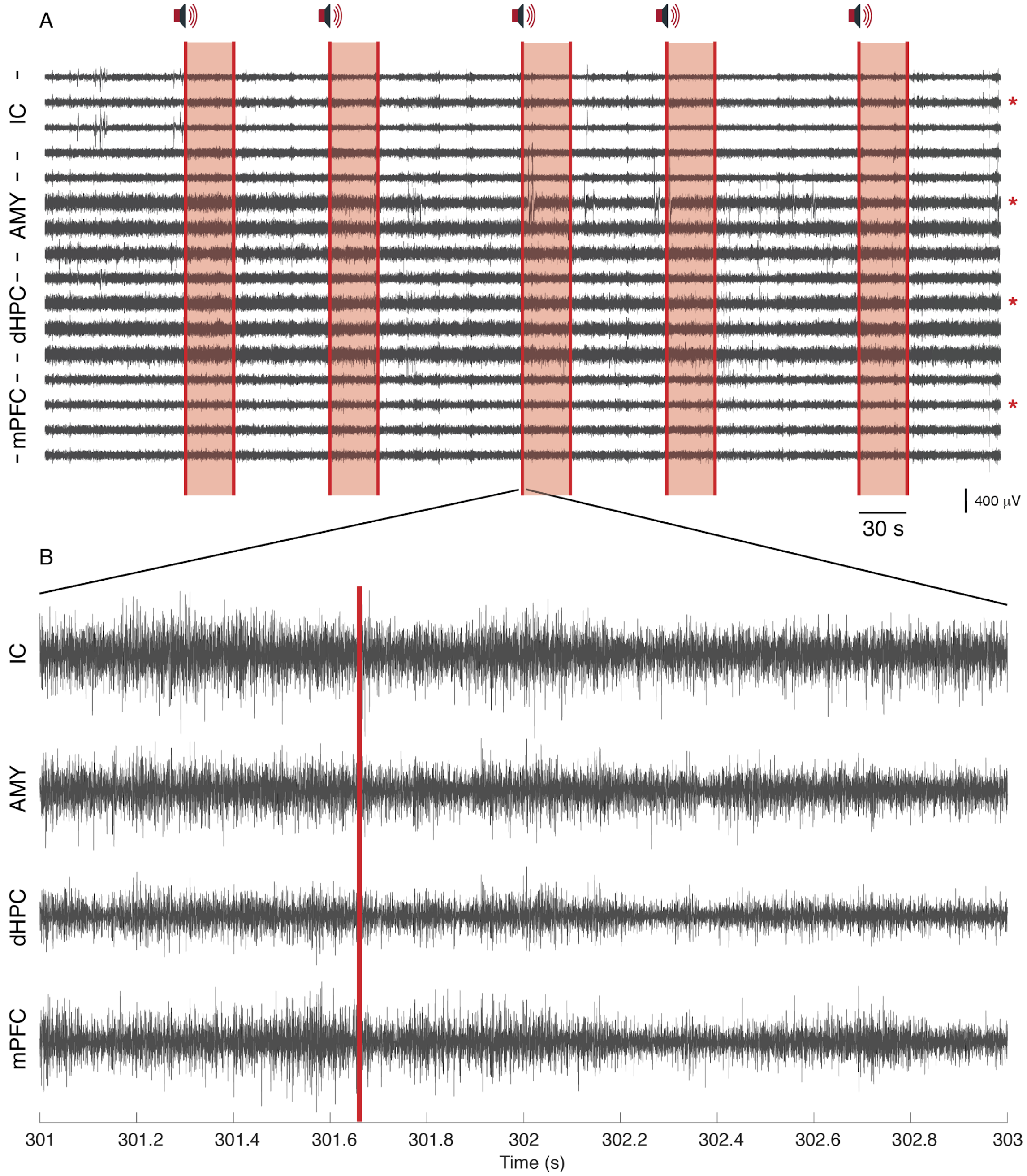


**Supplementary Figure 1. Single units from Mouse 1**. A) Raw record of sixteen channels on each brain substrate. In red we have the auditory stimulus presentation window. Each red asterisk represents the representative channels chosen. B) Two-second time window of the third auditory stimulus presentation. Filtered records between 300 and 3000 Hz.

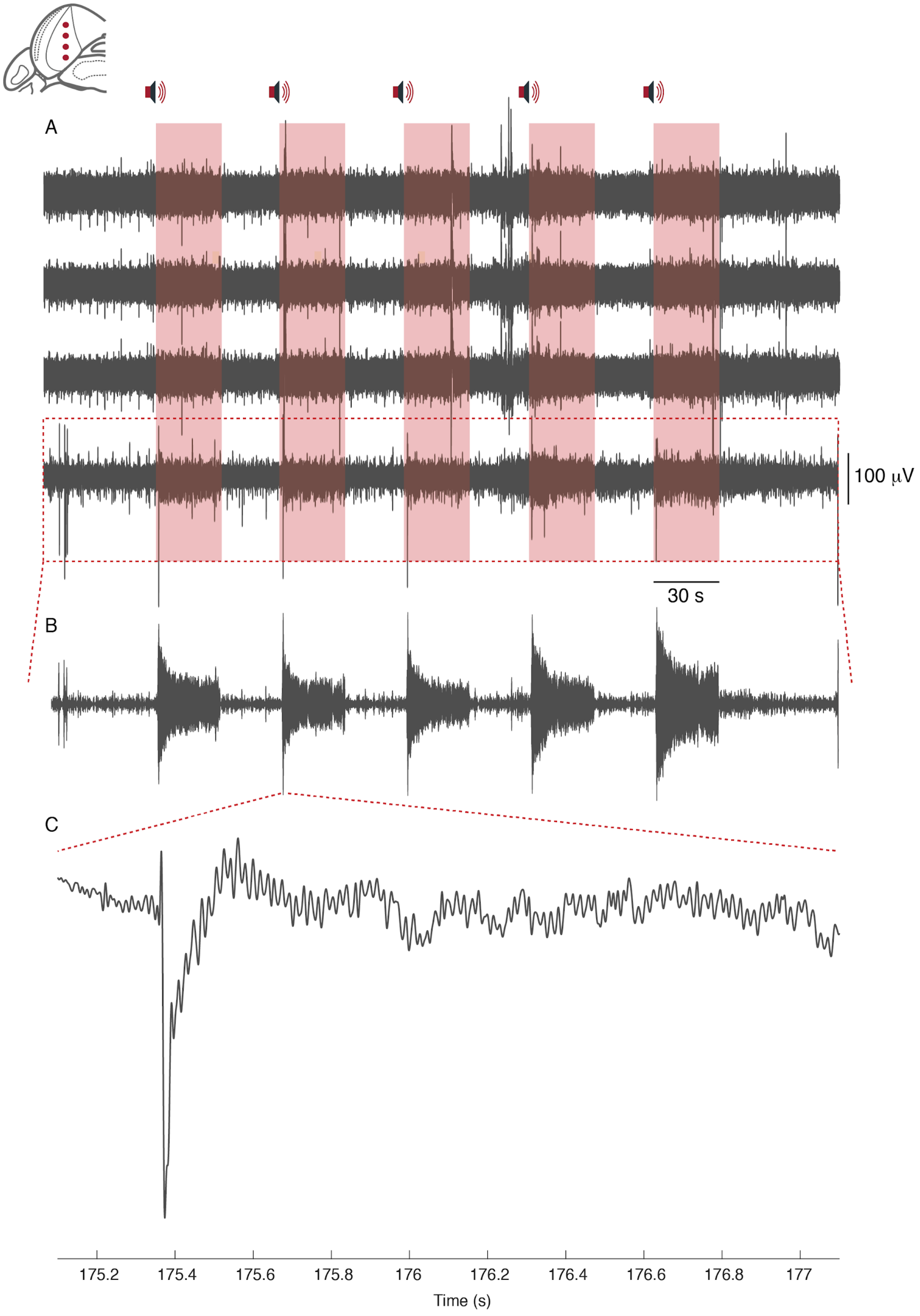


**Supplementary Figure 2. Local Field Potentials from Mouse 2**. A) Raw record of four channels in the IC. In red we have the auditory stimulus presentation window. B) Channel positioned in the ventral portion of the IC. Record filtered between 52 and 55 HZ. C) Two-second time window of the second auditory stimulus presentation. Filtered record between 1 and 100 Hz.
