## Supplementary for "A fully adapted headstage for electrophysiological experiments with custom and scalable electrode arrays to record widely distributed brain regions": Supplementary_Material.pdf

### Supplementary Material – Tutorials

**Authors:** Flávio Afonso Gonçalves Mourão[a], Leonardo de Oliveira Guarnieri [ab], Paulo Aparecido Amaral Júnior [ab], Vinícius Rezende Carvalho a[b], Eduardo Mazoni Andrade Marçal Mendes [ab], Márcio Flávio Dutra Moraes [a].

a. Núcleo de Neurociências, Departamento de Fisiologia e Biofísica, Instituto de Ciências Biológicas (ICB), Universidade Federal de Minas Gerais (UFMG). Av. Antônio Carlos, 6627 - CEP 31270-901. Belo Horizonte, Minas Gerais, Brazil.

b. Programa de Pós-Graduação em Engenharia Elétrica, Departamento de Engenharia Eletrônica (DELT), Escola de Engenharia, Universidade Federal de Minas Gerais (UFMG). Av. Antônio Carlos, 6627 - CEP 31270-901. Belo Horizonte, Minas Gerais, Brazil.

**Correspondence:**

Flávio Afonso Gonçalves Mourão, PhD


Márcio Flávio Dutra Moraes, PhD


2021

**Recording Headstage**

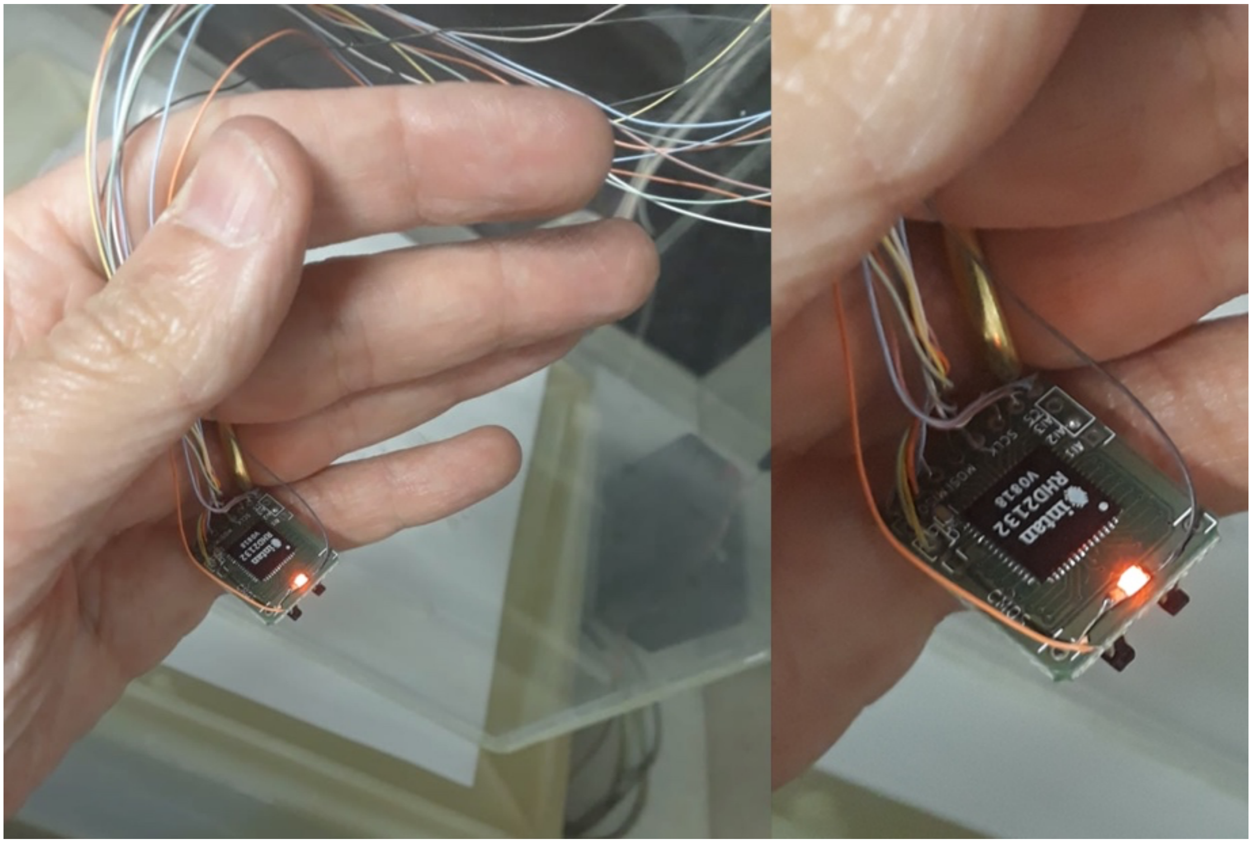


** Led is not in the original project*

| Component | Specification | Supplier | Price | Quantity | Link |
|---|---|---|---|---|---|
| FPC connector (Flexible Printed Circuits) | SMD vertical type 0.5mm pitch 2mm height | mktechnic | $0.79(un) | 2 | https://cutt.ly/Xx55qQE |
| RHD2132 amplifier chip | 32-channel unipolar inputs and common reference | Intan Technologies | $390 (un) | 1 | https://cutt.ly/Yx6qief |
| Resistor | SMD – 0 Ω | - | - | 1 | - |
| Capacitor | SMD – 10 nF | - | - | 1 | - |
| Capacitor | SMD – 100 nF | - | - | 2 | - |

### Components

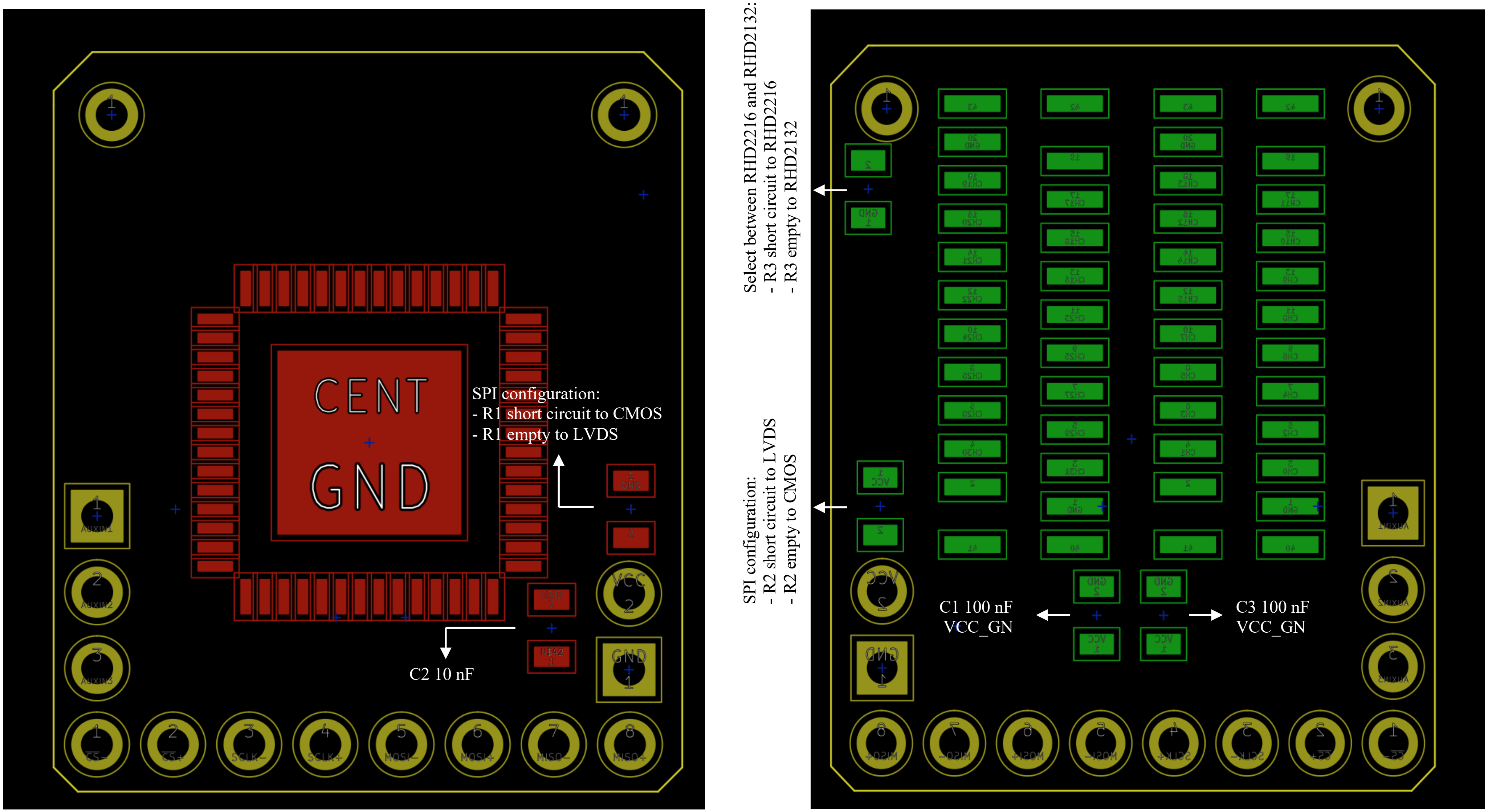
CENT
GND
SPI configuration:
- R1 short circuit to CMOS
- R1 empty to LVDS
C2 10 nF
Select between RHD2216 and RHD2132:
- R3 short circuit to RHD2216
- R3 empty to RHD2132
SPI configuration:
- R2 short circuit to LVDS
- R2 empty to CMOS
C1 100 nF
VCC_GN
C3 100 nF
VCC_GN

**Handmade cable**

We strongly recommend the following tutorial:

- https://open-ephys.atlassian.net/wiki/spaces/OEW/pages/491587/Fine+wire+tether

| Component | Specification | Supplier | Price | Quantity | Link |
|---|---|---|---|---|---|
| Omnetics Polarized PZN | Straight Thru-Hole (Type DD) 12 contacts | Omnetics Connector Corporation | *on request* | 1 | https://cutt.ly/FcqiCNQ |
| Micro bare copper wire | CZ 1187 wire series | Cooner Wire | *on request* | 12 | https://cutt.ly/lcqo9Pz |

**However, in our adapted board, each contact needs to be checked**

Wiring diagram:

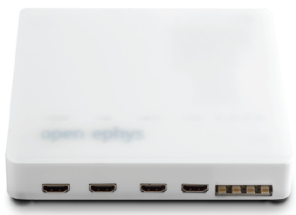

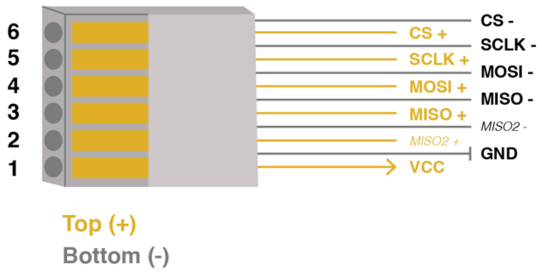


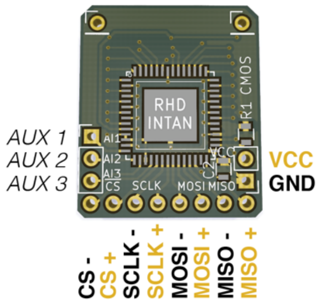


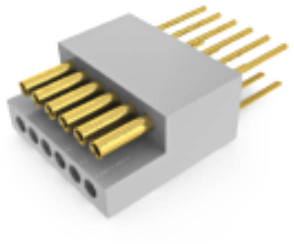

### Flat-Grid Connector – V2 – Current Version

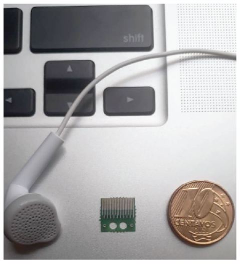


**Channels Map**

Be careful with the board position

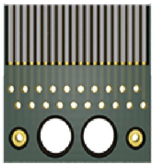

**Bottom view**

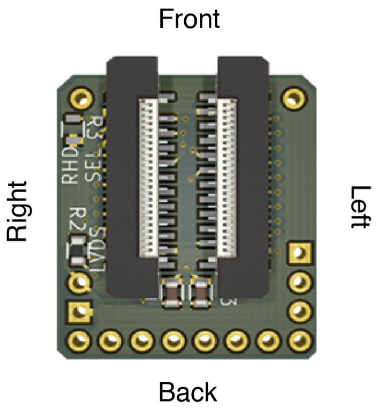


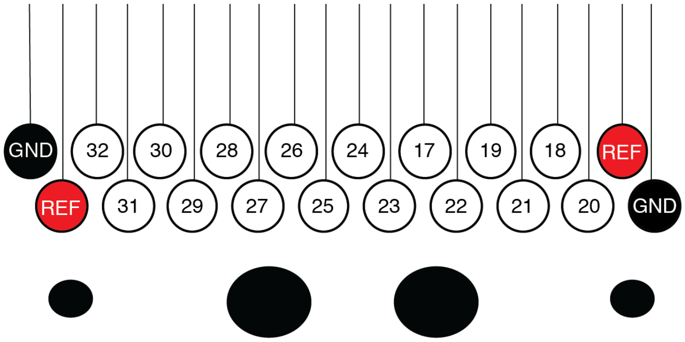


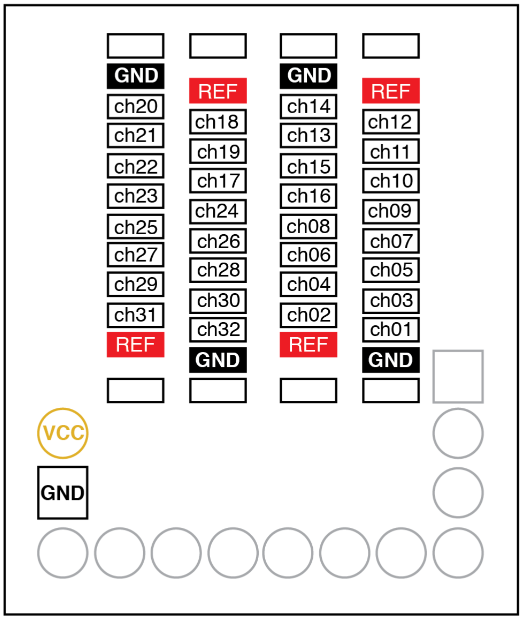


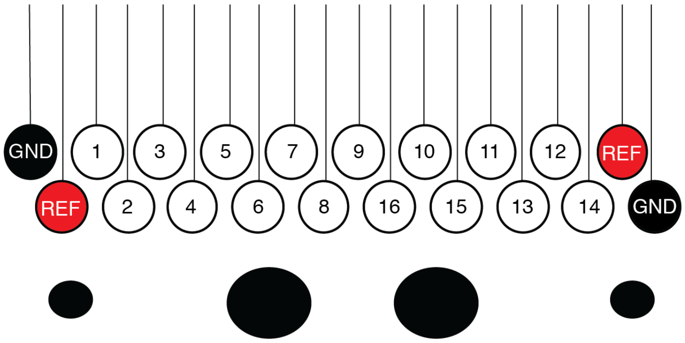


| Component | Specification | Supplier | Price | Quantity | Link |
|---|---|---|---|---|---|
| Stainless steel wire | 0.127 mm internal diameter, teflon-coated. 100 feet. Model 7914 | A-M Systems Inc | $140 (un) | 1 | https://cutt.ly/5x6eIoR |
| Tungsten microwires | | | | | |
| Silver paint | 1 troy oz. (31.1g) | SPI Supplies® | $50 (un) | 1 | https://cutt.ly/Ox6sup1 |
| Thermosetting polymer | 5 minute Epoxy | Devcon | - | | https://cutt.ly/Zx6ltrE<br>https://cutt.ly/mx6zobJ<br>https://cutt.ly/Rx6zFZZ |

### Flat-Grid Connector – V1 – Old Version

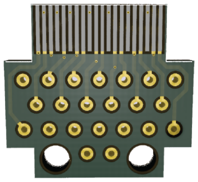

#### Channels Map

Be careful with the board position

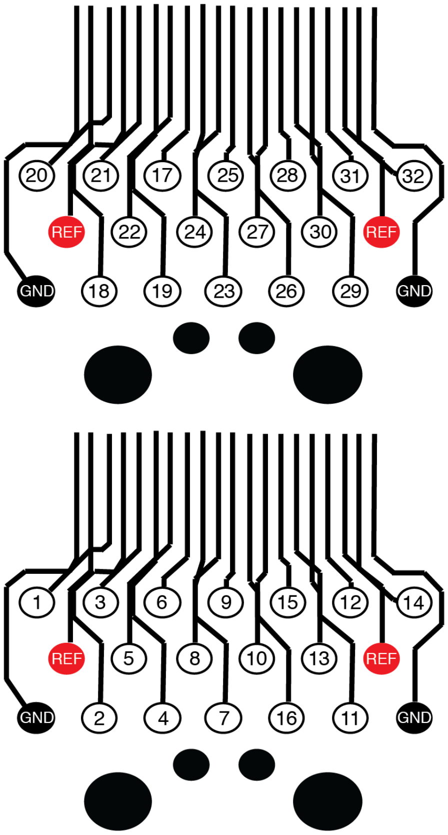


#### Bottom view

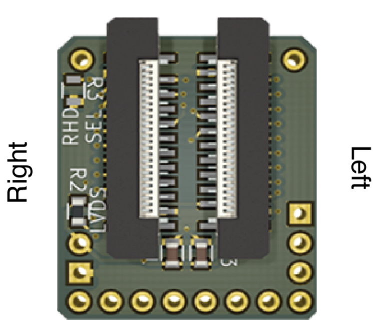


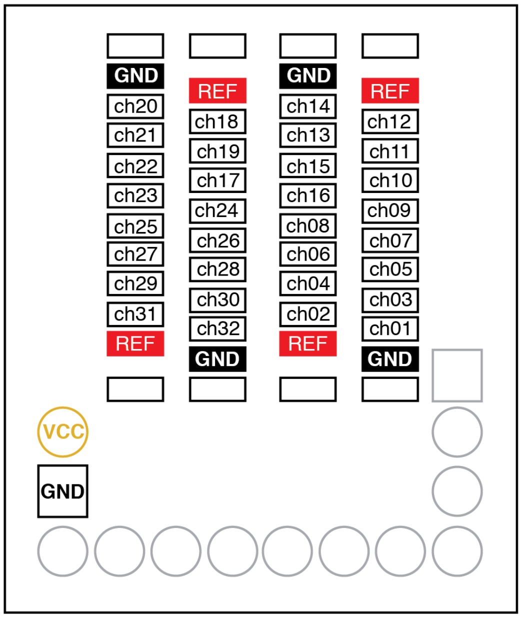

**SMD/FPC connector ← → Omnetics**
**Adapters for the original RHD headstage**

| Component | Specification | Supplier | Price | Quantity | Link |
|---|---|---|---|---|---|
| FPC connector (Flexible Printed Circuits) | SMD vertical type 0.5mm pitch 2mm height | mktechnic | $0.79(un) | 2 | https://cutt.ly/Xx55qQE |
| Omnetics Neuro Nano Strip | A79042-001 NPD-18-VV-GS | Omnetics Connector Corporation | *on request* | 1 | https://cutt.ly/UcqDO66 |
| Omnetics Neuro Nano Strip | A79046-001 NPD-36-DD-GS | Omnetics Connector Corporation | *on request* | 1 | https://cutt.ly/UcqDO66 |

**RHD 16-Channel Recording Headstages**

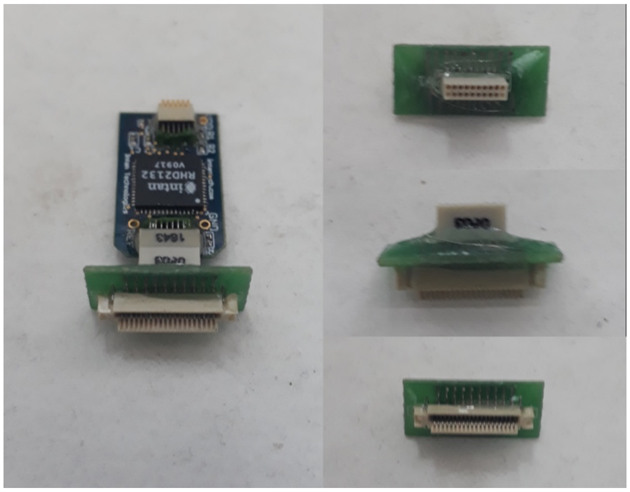

**RHD 32-Channel Recording Headstages**

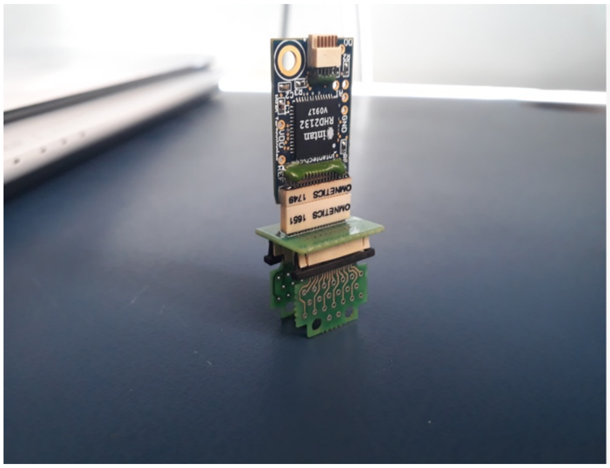
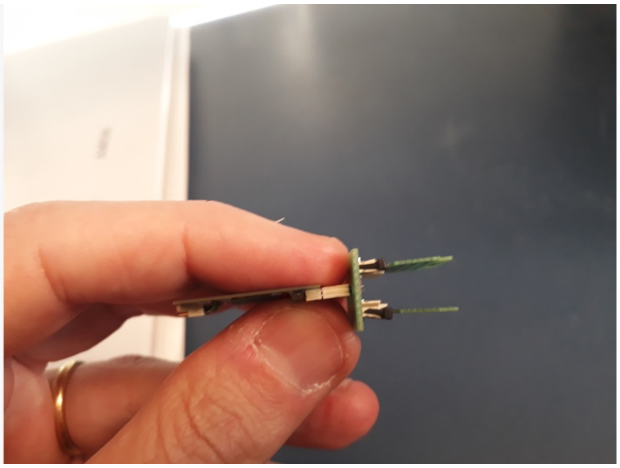
